# Chromosomal-level reference genome and microRNAs of the ricefield flatsedge *Cyperus iria*

**DOI:** 10.1101/2024.10.18.619046

**Authors:** Stacey S.K. Tsang, Wenyan Nong, Sean T.S. Law, Jacqueline C. Bede, Shan-Shan Chen, Sa Li, Yichun Xie, Thomas Swale, Yiqiang Zhao, David T.W. Lau, Zhen-peng Kai, Ting Fung Chan, Stephen S. Tobe, William G. Bendena, Jerome H.L. Hui

## Abstract

**Background:** Grass-like plants in the Cyperaceae family, commonly known as sedges, have a global distribution and include many economically problematic weeds. The ricefield flatsedge, *Cyperus iria*, is an aggressive weed in rice crops in Asia.

**Result:** Here, we present a chromosomal-level genome assembly for *C. iria* (461.2 Mb, scaffold N50 = 7.3 Mbp, 99.6% BUSCO score) providing potential targets for the control of this devastating weed. Based on the genome assembly and transcriptomes of vegetative tissues, 52,574 protein-coding genes were predicted to be encoded. A total of 26 conserved and 75 novel microRNAs, including 9 microRNA clusters, were also annotated. Synteny and microRNA cluster analyses further showed that *C. iria* had undergone at least one round of whole genome duplication.

**Conclusion:** The genomic resource established in this study sets up a foundation to further address basic and applied questions in the Cyperaceae.

## 1. Introduction

*Cyperus iria* belongs to the Cyperaceae, one of the largest monocot families with more than 5,400 described species [1]. *C. iria* is an annual, occasionally perennial, plant with a triangular culm with 3-ranked leaves [2](Fig.1A). The roots are reddish-brown fibrous roots without a rhizome. Given its rapid growth, prolific achene (seed) production (3,000-5,000 achenes/plant) and allelopathic specialized metabolites, *C. iria* is ranked among the top worst weeds, particularly in Asian rice crops where it can cause over 40% yield loss [3][4][5][6][7][8][9]. Thus, *C. iria*, which is widely distributed in its native range including Asia, Africa, and Australia, and now introduced across the Pacific Ocean to the United States (Fig. S1), is causing economically significant issues globally in agriculture and horticulture.

Part of the aggressive weedy nature of *C. iria* may be due to its allelopathic compounds such as sesquiterpenoid farnesol and juvenile hormone III (methyl-10R,11-epoxy-3,7,11-tri-methyl-2E,6E-dodecadienoate) that could have allelopathic effect on seeds germination on radish lettuce and rice [5]. Various studies also documented the inhibitory effect of *C. iria* on major crops such as different varieties of rice and soyabean [10][6][8]. Beside *C. iria*, allelopathic effect of other species from *Cyperus* genus was also reported, and biosynthesis of phenolic compounds was proposed to be the potential source of allelopathy [11]. A high-quality genome, thus, can provide fundamental information to study its adaptations to be a successful weed paddy field.

As well, *C. iria* is rich in flavonoids that act as antioxidants and is recognized as a medicinal plant [12]. *C. iria* extracts show hypoglycemic activity in streptozotocin-induced type 2 diabetic rats [13]. Volatile organic compounds released from *C. iria* leaves showed antifungal activity against *Fusarium graminearum* [14].

Despite the importance of *C. iria* as an aggressive weed and potential source of phytochemicals, currently only the chloroplast genome has been sequenced for this species [15]. To address this issue, here we provide a high quality chromosomal-level genome assembly for *C. iria* together with accompanying mRNA and small RNA transcriptomes.

## 2. Materials and Methods

### 2.1 Sample collections and DNA extraction and library preparation

Genomic DNA was extracted from a mature specimen of *Cyperus iria* collected in Sai Kung area of Hong Kong (Specimen No.: CUSLSH2052). Individuals for cultivation and transcriptomic studies were collected from the Sha Tin region of Hong Kong (Specimens No.: CUSLSH2163, CUSLSH2360 and CUSLSH2644). All specimens were deposited at the Shiu-Ying Hu Herbarium of The Chinese University of Hong Kong. Species identity was confirmed by sequencing the ITS1 and ITS2 regions. Heathy bracts, leaves and culms with no observable sign of pest damage, including bite marks, white waxy powder, crinkled leaves, and honeydew secretion or symptoms of pathogen infection including rust, mold, leaf spots, and chlorosis, were selected for DNA and RNA extraction. All plant material was extensively rinsed in sterile double-distilled water before DNA or RNA extraction. Genomic DNA was extracted using cetyltrimethylammonium bromide (CTAB) with the addition of 1% polyvinylpyrrolidone (PVP)[16]. The eluted DNA was subjected to quality control using a NanoDrop™ One Microvolume UV-Vis Spectrophotometer (Thermo Scientific), Qubit^®^ Fluorometer and overnight pulse-field gel electrophoresis. DNA hydrodynamic shearing was performed by transferring diluted DNA (40 ng/μL) into a g-TUBE (Covaris 520079) followed by 6 passes of centrifugations at 2,000 × g for 2 minutes. Further purification of sheared DNA was carried out using SMRTbell^®^ cleanup beads (PacBio Ref. No. 102158-300). After overnight pulse-field gel electrophoresis, the library was constructed using the SMRTbell® prep kit 3.0 (PacBio Ref. No. 102-141-700) following the manufacturer’s protocol. The final library preparation for sequencing was conducted using The Sequel^®^ II binding kit 3.2 (PacBio Ref. No. 102-194-100) The library was loaded on-plate in diffusion mode at concentration of 90 pM followed by sequencing on the Pacific Biosciences SEQUEL IIe System at The Chinese University of Hong Kong under the following conditions: run 30 hour with 2 hour pre-extension to generate HiFi reads. Details of the sequencing data can be found in Table S2A.

For each Dovetail Omni-C library, chromatin was fixed in place with formaldehyde in the nucleus and then extracted. Fixed chromatin was digested with DNAse I, chromatin ends were repaired and ligated to a biotinylated bridge adapter followed by proximity ligation of adapter containing ends. After proximity ligation, crosslinks were reversed and the DNA purified. Purified DNA was treated to remove biotin that was not internal to ligated fragments. Sequencing libraries were generated using NEBNext Ultra enzymes and Illumina-compatible adapters. Biotin-containing fragments were isolated using streptavidin beads before PCR enrichment of each library. The library was sequenced on an Illumina HiSeqX platform at the Dovetail Genomics to produce 190 million 2 × 150 bp paired end reads. Details of the sequencing data can be found in Table S2A.

### 2.2 Total and small RNA sequencing

All samples of different tissues were pretreated with CTAB [17], total RNA of samples CyI_K(1) and CyI_L(1) were extracted with TRIzol and the other samples used mirVana miRNA Isolation Kit (Ambion) following the manufacturer’s instructions. The extracted total RNA was subjected to quality and quantity analyses by gel electrophoresis, Nanodrop spectrophotometer (ThermoScientific) and a 2100 Bioanalyser (Agilent). Qualified total RNA samples were further sent to Novogene for library construction and sequencing.

After quality control, mRNA was enriched using oligo(dT) beads and rRNA was removed using the Ribo-Zero kit. The mRNA was fragmented randomly with the use of fragmentation buffer, then the cDNA is synthesized based on the mRNA template with random hexamers primer. The second-strand synthesis was then proceeded by adding a custom second-strand synthesis buffer (Illumina), dNTPs, RNase H and DNA polymerase I. After terminal repair, ligation and addition of sequencing adaptors, the double-stranded cDNA library was completed through size selection and PCR enrichment. Insert sizes and library concentrations of final libraries were determined using the 2100 Bioanalyser (Agilent) and real-time quantitative PCR (TaqMan Probe) respectively. The qualified libraries were pooled according to concentration and sequenced (Illumina) (Table S2B).

For Small RNA sequencing, aliquots of total RNA samples were submitted to Novogene for HiSeq Small RNA library construction and 50 bp single-end sequencing. A library was constructed by Small RNA Sample Pre Kit for the qualified samples, namely CyI_L3_2s, CyI_L1_3s, CyI_R3_3s and CyI_L1_As (Table S2B) which passed the quality checking as previously mentioned. The final cDNA library was ready for sequencing after a round of sequencing adaptor ligation, reverse transcription, PCR enrichment, purification and size selection.

### 2.3 Genome assembly

*De novo* genome assembly was performed using Hifiasm [18], which was then searched against the NT database using BLAST for the input for BlobTools (v1.1.1) to identify and remove any possible contaminations [19] (Table S2A). Haplotypic duplications were removed using “purge_dups” based on the depth of HiFi reads [20]. Furthermore, proximity ligation data sequenced from the Omni-C library were employed to scaffold the assembly with YaHS [21]. k-mers of the Omni-C reads were counted using jellyfish version 2.2.10 [22] with k = 31 and estimation of genome size, repeat content, and heterozygosity were analyzed based on a k-mer-based statistical approach using the GenomeScope webtool (http://genomescope.org/genomescope2.0/) [23].

### 2.4 Gene model prediction

RNA sequencing reads was pre-processed with trimmomatic (v0.33) with parameters “ILLUMINACLIP:TruSeq3-PE.fa:2:30:10 SLIDINGWINDOW:4:5 LEADING:5 TRAILING:5 MINLEN:25” [24] and cleaned with Kraken 2 to remove the contamination (kraken database k2_standard_20210517) [25]. The genome was soft-masked using redmask (v0.0.2). Genome annotation was performed using funannotate (v1.8.5, https://github.com/nextgenusfs/funannotate) [26]. Briefly, the processed RNA reads and the softmasked genome were used to run “funannotate train” with parameters “--stranded RF -- max_intronlen 350,000” to align RNA-seq data, ran Trinity, and then ran PASA [27]. The PASA gene models were used to train Augustus in “funannotate predict” step following recommended options for eukaryotic genomes (https://funannotate.readthedocs.io/en/latest/tutorials.html#non-fungal-genomes-higher-eukaryotes). Briefly, the gene models were predicted by funannotate predict using the following parameters: “--repeat-s2evm --protein_evidence uniprot_sprot.fasta -- genemark_mode ET --busco_seed_species arabidopsis --optimize_augustus --busco_db embryophyta --organism other --max_intronlen 350,000”. The gene models originated from several prediction sources, including: GeneMark [28] HiQ, pasa, Augustus [29], GlimmerHMM [30], snap [31]. Gene models were passed to Evidence Modeler [27] (EVM Weights: [GeneMark: 1, HiQ: 2, pasa: 6, proteins: 1, Augustus: 1, GlimmerHMM: 1, snap: 1, transcripts: 1]) to generate the final annotation files, and PASA [27] was used to update the EVM consensus predictions, add UTR annotations and models for alternatively spliced isoforms.

### 2.5 Repetitive elements annotation

Annotation of transposable elements (TEs) was conducted using the Earl Grey TE annotation workflow pipeline (version 1.2, https://github.com/TobyBaril/EarlGrey) [32].

### 2.6 microRNA annotation

To process small RNA data, small RNA sequencing raw reads with Phred quality score less than 20 were removed and adaptor sequences were trimmed. MicroRNA annotation was performed using sRNAminer (v1.1.2)[33]. Briefly, processed reads were collapsed by Seq_collapsing and rRNA, tRNA, snoRNA, snRNA and organelle fragments were removed by Data_cleaning using the ncRNA database Rfam_withoutMIR.fa and organelle database Organelle.genomic.fa downloaded from sRNAminer databases, allowing 1 mismatch in alignment and 20 of the locations a read can map to. The microRNA was identified using miRNA_identification_no_align with parameters 20 and 22 of the minimum and maximum length of the miRNA and 300 of the maximum length of the miRNA loop. Digital expression of microRNA expression was calculated by miRNA_abundance, the microRNA with abundance RP10M (reads per 10 million) > 1 was marked as a microRNA.

## 3. Results and Discussion

### 3.1 High-quality *Cyperus iria* genome

The *Cyperus iria* genome was sequenced and assembled using PacBio HiFi reads followed by further scaffolding with Omni-C reads (Table S1). The genome assembly was determined to be 461.2 Mbp with a high sequence contiguity of scaffold N50 = 7.3 Mbp and high sequence completeness with a 99.6% complete BUSCO score for viridiplantae genes (version odb10) (Fig. 1B). Most of the sequences (∼99.9%) were anchored to 69 pseudomolecules (Fig. 1C, Table S3), and a total of 49,202 gene models including 780 tRNA genes and 52,574 protein-coding gene were predicted using the transcriptomes generated in this study. A total of 156.75 Mb repetitive elements were also annotated, accounting for 33.98 % of the genome assembly. DNA transposons accounted for 3.79% of the genome, while long terminal repeat (LTR), long interspersed nuclear elements (LINE) and short interspersed nuclear elements (SINE) belonging to retrotransposons accounted for 7.87%, 0.80% and 0.001% of the genome, respectively Table S4).

**Figure 1.**
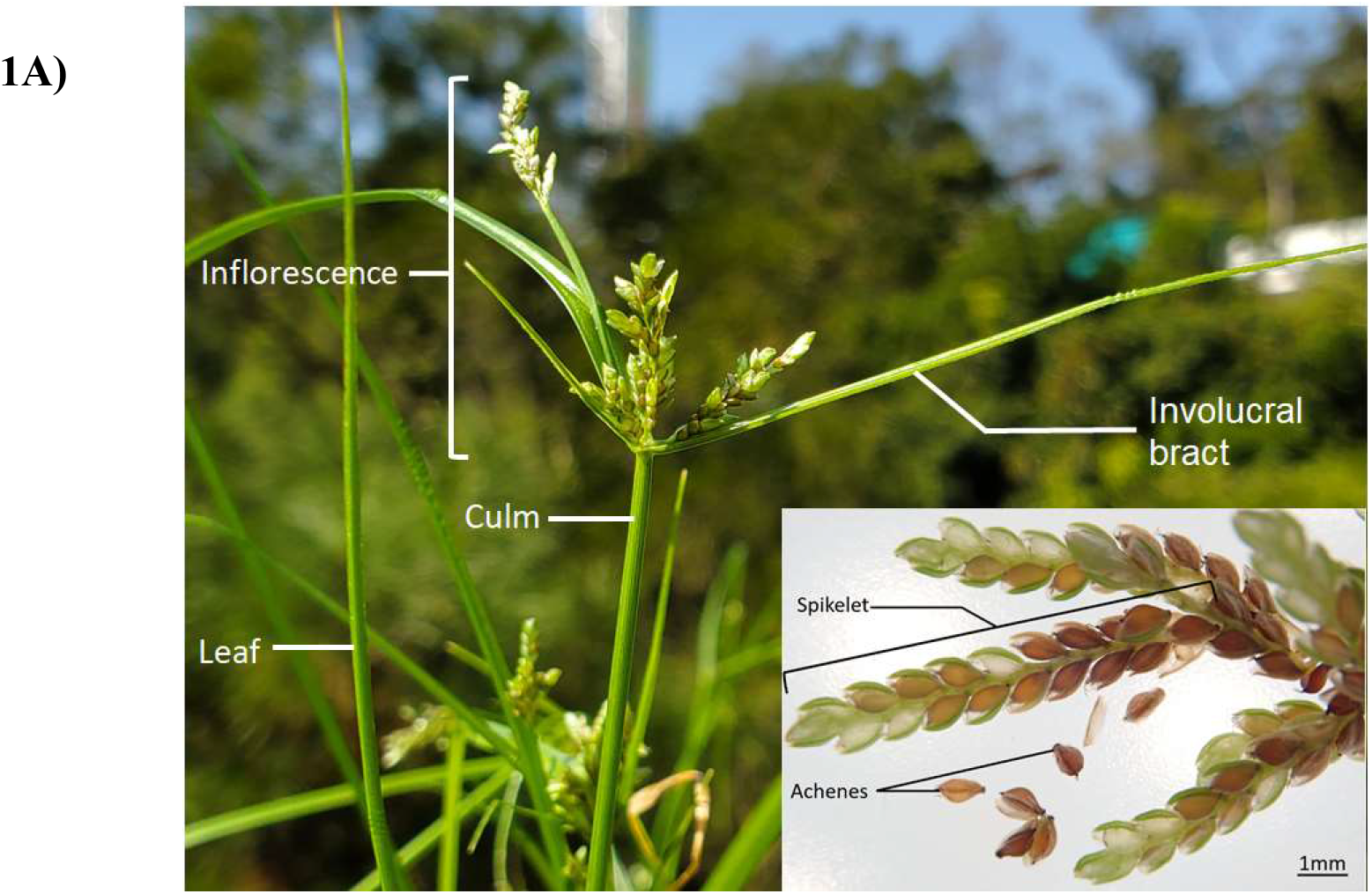
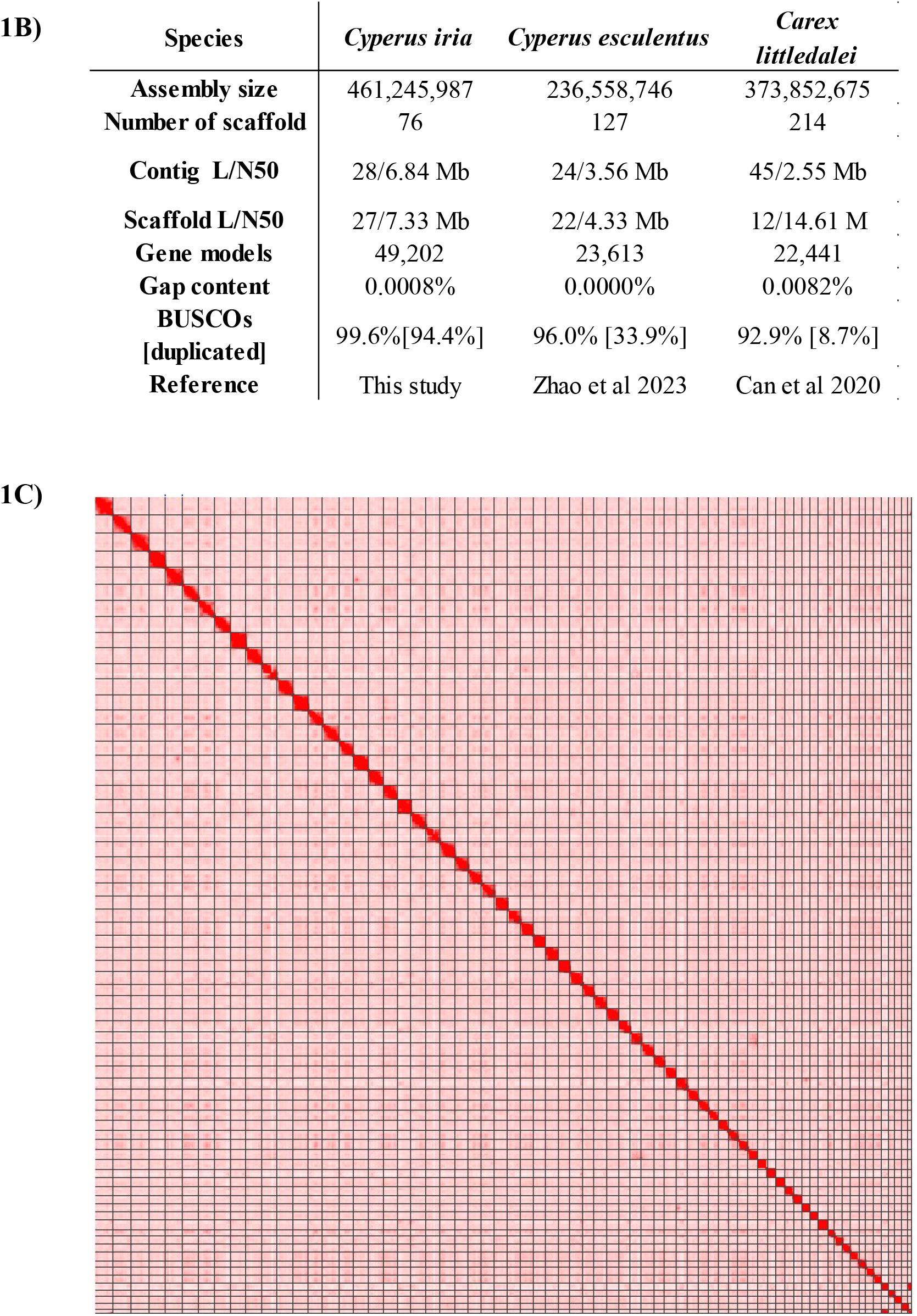
A) Picture showing the morphology of *Cyperus iria*; B) Genome statistics; C) High-throughput chromatin conformation capture (Omni-C) interaction map of *Cyperus iria* visualizes the number of chromosome interactions within and between 69 chromosomes.

### 3.2. MicroRNAs

MicroRNAs impact plant survival in both abiotic and biotic stresses [34][35]. However, our knowledge on microRNAs in the Cyperaceae is currently lacking. A total of 239 microRNAs were identified, including 109 known, 75 novel microRNAs, and 9 potential microRNA clusters visualized in Fig. 2 with TBtools [36] (Tables S6-7). By comparison to other published plant genomes, 21 out of the 23 miRNA families are found to be conserved among all angiosperms, while miR-2275 and miR-528 families are only conserved between members of monocotyledons. miR-2275 has been found to be associated with nitrogen acquisition enhancement in wheat which allow the plant to adapt nitrogen limited situation [37]. miR-528 was found to target genes that involved in reactive oxygen species (ROS) metabolism which is necessary in responses to extreme temperature [38]. It is worth noting that miR-1432, which was previously only identified in the Poaceae family, is now also identified in *C. iria*. miR-1432 has previously been demonstrated to contribute to disease resistance and drought stress tolerance in rice [39], [40]. The identification of specific microRNAs in *Cyperus* sets up a foundation for further investigation of their roles in the Cyperaceae.

**Figure 2.**
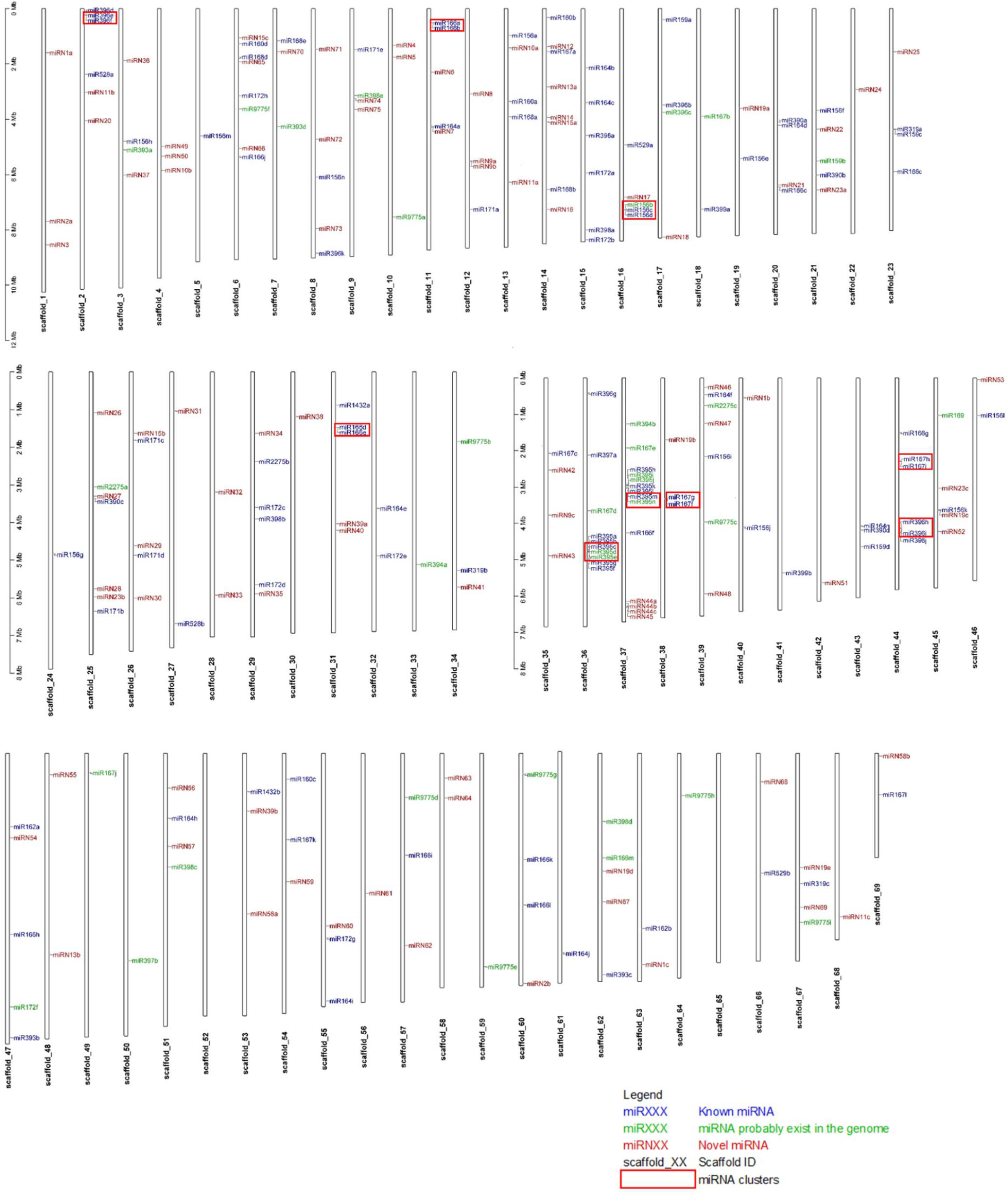
Genomic loci of microRNAs in *C. iria*.

### 3.3. Whole genome duplication

Whole genome duplication (WGD) events sigma *ρ* and rho *σ* are closely associated with the origin of grass family [41] that occurred during Cretaceous and Paleogene periods. Recent genomic studies on the Cyperaceae revealed different WGD events occurred in different clades. For instance, there was no large-scale genome duplication inferred in the genomes of *Kobresia myosuroides* and *Bolboschoenus planiculmis* [42][43], while a clade-specific WGD occurred after the divergence of Schoenoplecteae and Bolboschoeneae [44], as well as after the divergence of *Carex littledalei* and *Rhynchospora breviuscula* [45][46][47]. In the syntenic analyses between the longest 69 genomic scaffolds of *C. iria*, all of them had syntenic relationships with at least another scaffold in the genome assembly which was visualized with TBtools in Fig. 3A [36]. Given microRNAs are also good indicators for whole genome duplication events [48][49], we have also performed syntenic analyses and revealed that all microRNA clusters had a 1-to-1 syntenic relationship suggesting that at least one round of whole genome duplication happened in the *C. iria* genome (Fig. 3B).

**Figure 3.**
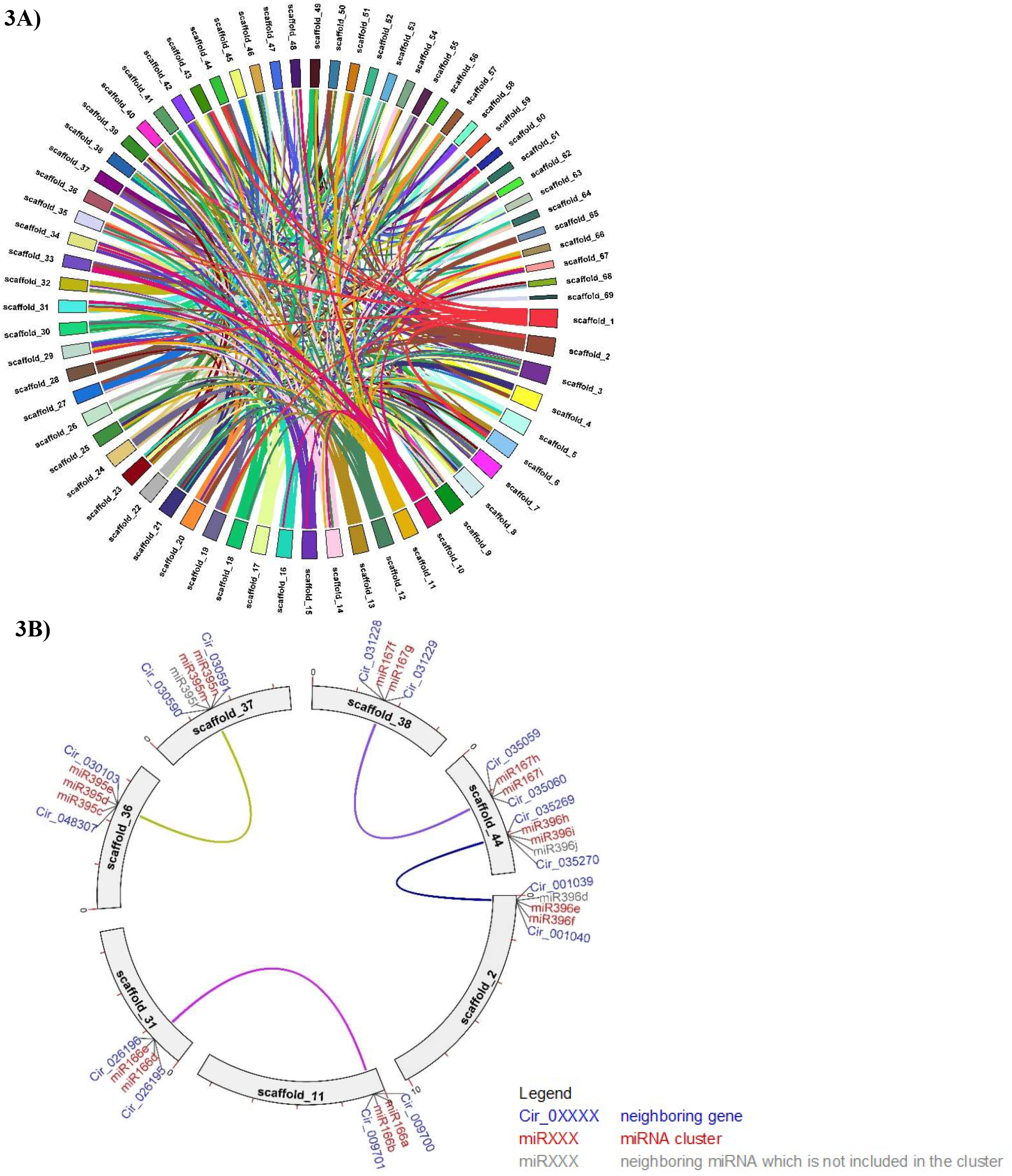
**A)** Self-syntenic analysis of *C. iria* genome assembly; **B)** Synteny analysis of microRNA clusters and their neighboring genes.

## 4. Conclusion

This study presents the first high-quality genome and transcriptomic resources for *Cyperus iria*. In addition, we have annotated microRNAs and syntenic analyses suggested that at least one round of whole genome duplication happened in this species. Together, this study provides a foundation for further study of Cyperaceae biology and evolution.

## Data availability

The raw reads generated in this study have been deposited to the NCBI database under the BioProject accession number PRJNA681037. The genome assemblies and genome annotation files were deposited in Figshare (https://doi.org/10.6084/m9.figshare.19235748).

## Acknowledgements

This work was supported by State Key Laboratory of Agrobiotechnology (8300040), Hong Kong Research Grant Council Collaborative Research Fund (C4015-20EF), General Research Fund (14100420), and Direct Grants of The Chinese University of Hong Kong (4053489). SSKT was supported by the PhD studentship provided by The Chinese University of Hong Kong.

## References

[1] Govaerts R, Simpson DA, World Checklist of Cyperaceae. Sedges. London: The Board of Trustees of the Royal Botanic Gardens, Kew., 2007.

[2] Deng YF, Flora of Hong Kong Volume 4. Hong Kong: Agriculture, Fisheries and Conservation Department, 2011.

[3] Holm LG, Plucknett DL, Pancho JV, and Herberger JP, The world’s worst weeds. Distribution and biology. Honolulu, Hawaii, USA: University Press of Hawaii, 1977.

[4] Ampong-Nyarko K, De Datta SK, A Handbook for Weed Control in Rice. Manila, Philippines: Intermational Rice Research Institute, 1991.

[5] Bede JC and Tobe SS, Activity of insect juvenile hormone III: Seed germination and seedling growth studies. Chemoecology, 2000; 10(2): 89–97, 2000, doi: 10.1007/s000490050012.

[6] Ismail BS, Siddique MA. The Inhibitory Effect of Grasshopper’s Cyperus (Cyperus iria L.) on the Seedling Growth of Five Malaysian Rice Varieties. Trop Life Sci Res. 2011; 22(1):81–9.

[7] Dhammu, HS, Sandhu, KS Critical period of Cyperus iria L. competition in transplanted rice. In Proceedings of the 13th Aust. Weeds Conf.: Weeds “Threats now and forever?”, Perth, Australia, 2002; 79–82.

[8] Chopra N, Tewari G, Tewari L, Upreti B, Pandey N. Allelopathic Effect of Echinochloa colona L. and Cyperus iria L. Weed Extracts on the Seed Germination and Seedling Growth of Rice and Soyabean. Adv. Agric. 2017; 2017(4):1–5. doi: 10.1155/2017/5748524.

[9] Awan TH, Ali HH, Chauhan BS. Cyperus iria Weed Growth, Survival, and Fecundity in Response to Varying Weed Emergence Times and Densities in Dry-Seeded Rice Systems. Agronomy. 2022; 12(5):1–15. doi:10.3390/agronomy12051006.

[10] Manandhar S, Shrestha BB, Lekhak HD, “Weeds of Paddy Field at Kirtipur, Kathmandu,” Sci. World, 2007; 5(5):100–106, doi: 10.3126/sw.v5i5.2665.

[11] Silva A, Bezerra JJ, Prata A, Souza R, Paulino C, Nascimento T, Silva S, Costa V, Duarte M. Phytochemical Profile and Evaluation of the Allopathic Effect of Three Species of the Genus Cyperus (Cyperaceae). J. Agric. Stud. 2020, 8 (3): 569–584, doi: 10.5296/jas.v8i3.16724.

[12] Ho YL, Huang SS, Deng JS, Lin YH, Chang YS, Huang GJ. In vitro antioxidant properties and total phenolic contents of wetland medicinal plants in Taiwan. Bot. Stud. 2012; 53(1):55–66.

[13] Saeed M, Sharif A, Hassan SU, Akhtar B, Muhammad F, Malik M. Cyperus iria aqueous-ethanol extract ameliorated hyperglycemia, oxidative stress, and regulated inflammatory cytokines in streptozotocin-induced diabetic rats. Environ Sci Pollut Res Int. 2022; 29(3):4769–4784. doi: 10.1007/s11356-021-15917-9.

[14] Jiang Y, Ownley BH, Chen F. Terpenoids from Weedy Ricefield Flatsedge (Cyperus iria L.) Are Developmentally Regulated and Stress-Induced, and have Antifungal Properties. Molecules. 2018; 23(12):3149. doi: 10.3390/molecules23123149

[15] Yang LL, Niu JQ, Tang WW. The complete chloroplast genome of pioneering plant Cyperus iria L. (Cyperaceae) in ecological restoration. Mitochondrial DNA B Resour. 2021; 6(4):1335–1336. doi: 10.1080/23802359.2021.1908865.

[16] Doyle JJ, Doyle JL. A Rapid DNA Isolation Procedure from Small Quantities of Fresh Leaf Tissues. Phytochem Bull. 1986; 19(1):11–15.

[17] Jordon-Thaden IE, Chanderbali AS, Gitzendanner MA, Soltis DE. Modified CTAB and TRIzol protocols improve RNA extraction from chemically complex Embryophyta. Appl Plant Sci. 2015;3(5):1400105. doi: 10.3732/apps.1400105.

[18] Cheng H, Concepcion GT, Feng X, Zhang H, Li H. Haplotype-resolved de novo assembly using phased assembly graphs with hifiasm. Nat Methods. 2021;18(2):170–175. doi: 10.1038/s41592-020-01056-5.

[19] Laetsch DR, Blaxter ML, Leggett RM, “BlobTools : Interrogation of genome assemblies,” F1000 Res. 2017; 6(1287):1–16.

[20] Guan D, McCarthy SA, Wood J, Howe K, Wang Y, Durbin R. Identifying and removing haplotypic duplication in primary genome assemblies. Bioinformatics. 2020; 36(9):2896–2898. doi: 10.1093/bioinformatics/btaa025.

[21] Zhou C, McCarthy SA, Durbin R. YaHS: yet another Hi-C scaffolding tool. Bioinformatics. 2023; 39(1):1–3. doi: 10.1093/bioinformatics/btac808.

[22] Marçais G, Kingsford C. A fast, lock-free approach for efficient parallel counting of occurrences of k-mers. Bioinformatics. 2011; 27(6):764–770. doi: 10.1093/bioinformatics/btr011.

[23] Vurture GW, Sedlazeck FJ, Nattestad M, Underwood CJ, Fang H, Gurtowski J, Schatz MC. GenomeScope: fast reference-free genome profiling from short reads. Bioinformatics. 2017; 33(14):2202–2204. doi: 10.1093/bioinformatics/btx153.

[24] Bolger AM, Lohse M, Usadel B. Trimmomatic: a flexible trimmer for Illumina sequence data. Bioinformatics. 2014; 30(15):2114–2120. doi: 10.1093/bioinformatics/btu170.

[25] Wood DE, Salzberg SL. Kraken: ultrafast metagenomic sequence classification using exact alignments. Genome Biol. 2014;15(3):R46. doi: 10.1186/gb-2014-15-3-r46.

[26] Palmer J, Stajich JE, Funannotate: Eukaryotic genome annotation pipeline. (2017). https://funannotate.readthedocs.io/en/latest/annotate.html (accessed Feb. 05, 2022).

[27] Haas BJ, Salzberg SL, Zhu W, Pertea M, Allen JE, Orvis J, White O, Buell CR, Wortman JR. Automated eukaryotic gene structure annotation using EVidenceModeler and the Program to Assemble Spliced Alignments. Genome Biol. 2008; 9(1):R7. doi: 10.1186/gb-2008-9-1-r7.

[28] Lomsadze A, Ter-Hovhannisyan V, Chernoff YO, Borodovsky M. Gene identification in novel eukaryotic genomes by self-training algorithm. Nucleic Acids Res. 2005; 33(20):6494–6506. doi: 10.1093/nar/gki937.

[29] Stanke M, Keller O, Gunduz I, Hayes A, Waack S, Morgenstern B. AUGUSTUS: ab initio prediction of alternative transcripts. Nucleic Acids Res. 2006; 34(Web Server issue):W435–9. doi: 10.1093/nar/gkl200.

[30] Majoros WH, Pertea M, Antonescu C, Salzberg SL. GlimmerM, Exonomy and Unveil: three ab initio eukaryotic genefinders. Nucleic Acids Res. 2003; 31(13):3601–3604. doi: 10.1093/nar/gkg527.

[31] Korf I. Gene finding in novel genomes. BMC Bioinformatics. 2004; 5(1):1–9. doi: 10.1186/1471-2105-5-59.

[32] Baril T, Galbraith J, Hayward A. Earl Grey: A Fully Automated User-Friendly Transposable Element Annotation and Analysis Pipeline. Mol Biol Evol. 2024; 41(4):msae068. doi: 10.1093/molbev/msae068.

[33] Li G, Chen C, Chen P, Meyers BC, Xia R. sRNAminer: A multifunctional toolkit for next-generation sequencing small RNA data mining in plants. Sci Bull (Beijing). 2024; 69(6):784–791. doi: 10.1016/j.scib.2023.12.049.

[34] Liu SR, Zhou JJ, Hu CG, Wei CL, Zhang JZ. MicroRNA-Mediated Gene Silencing in Plant Defense and Viral Counter-Defense. Front Microbiol. 2017; 8:1801. doi: 10.3389/fmicb.2017.01801.

[35] Pérez-Quintero Á.L, Neme R, Zapata A, López C. Plant microRNAs and their role in defense against viruses: a bioinformatics approach. BMC Plant Biol. 2010; 10 (1):1–12. doi: 10.1186/1471-2229-10-138.

[36] Chen C, Chen H, Zhang Y, Thomas HR, Frank MH, He Y, Xia R. TBtools: An Integrative Toolkit Developed for Interactive Analyses of Big Biological Data. Mol Plant. 2020; 13(8):1194–1202. doi: 10.1016/j.molp.2020.06.009

[37] Qiao Q, Wang X, Yang M, Zhao Y, Gu J, Xiao, K. Wheat miRNA member TaMIR2275 involves plant nitrogen starvation adaptation via enhancement of the N acquisition-associated process, Acta Physiol. Plant. 2018; 40(183):1–13 doi: 10.1007/s11738-018-2758-9.

[38] Zhu H, Chen C, Zeng J, Yun Z, Liu Y, Qu H, Jiang Y, Duan X, Xia R. MicroRNA528, a hub regulator modulating ROS homeostasis via targeting of a diverse set of genes encoding copper-containing proteins in monocots. New Phytol. 2020; 225(1):385–399. doi: 10.1111/nph.16130.

[39] Li Y, Zheng YP, Zhou XH, Yang XM, He XR, Feng Q, Zhu Y, Li GB, Wang H, Zhao JH, et al. Rice miR1432 Fine-Tunes the Balance of Yield and Blast Disease Resistance via Different Modules. Rice (N Y). 2021; 14(1):87. doi: 10.1186/s12284-021-00529-1.

[40] Luo G, Li L, Yang X, Yu Y, Gao L, Mo B, Chen X, Liu L. MicroRNA1432 regulates rice drought stress tolerance by targeting the CALMODULIN-LIKE2 gene. Plant Physiol. 2024;195(3):1954–1968. doi: 10.1093/plphys/kiae127.

[41] Paterson AH, Bowers JE, Chapman BA. Ancient polyploidization predating divergence of the cereals, and its consequences for comparative genomics. Proc Natl Acad Sci U S A. 2004; 101(26):9903–9908. doi: 10.1073/pnas.0307901101.

[42] Ning Y, Li Y, Dong SB, Yang HG, Li CY, Xiong B, Yang J, Hu YK, Mu XY, Xia XF. The chromosome-scale genome of Kobresia myosuroides sheds light on karyotype evolution and recent diversification of a dominant herb group on the Qinghai-Tibet Plateau. DNA Res. 2023; 30(1):dsac049. doi: 10.1093/dnares/dsac049.

[43] Ning Y, Li Y, Lin HY, Kang EZ, Zhao YX, Dong SB, Li Y, Xia XF, Wang YF, Li CY. Chromosome-Scale Genome Assembly for Clubrush (Bolboschoenus planiculmis) Indicates a Karyotype with High Chromosome Number and Heterogeneous Centromere Distribution. Genome Biol Evol. 2024;16(3):evae039. doi: 10.1093/gbe/evae039.

[44] Li Y, Ning Y, Zheng YC, Lou XY, Pan Z, Dong SB. Chromosome-Scale Genome Assembly for Soft-Stem Bulrush (Schoenoplectus tabernaemontani) Confirms a Clade-Specific Whole-Genome Duplication in Cyperaceae. Genome Biol Evol. 2024; 16(7):1–8. doi: 10.1093/gbe/evae141.

[45] Zhao X, Yi L, Ren Y, Li J, Ren W, Hou Z, Su S, Wang J, Zhang Y, Dong Q, Yang X, Cheng Y, Lu Z. Chromosome-scale Genome Assembly of the Yellow Nutsedge (Cyperus esculentus). Genome Biol Evol. 2023;15(3):1–7. doi: 10.1093/gbe/evad027.

[46] Hofstatter PG, Thangavel G, Lux T, Neumann P, Vondrak T, Novak P, Zhang M, Costa L, Castellani M, Scott A, et al. Repeat-based holocentromeres influence genome architecture and karyotype evolution. Cell. 2022;185(17):3153-3168.e18. doi: 10.1016/j.cell.2022.06.045.

[47] Can M, Wei W, Zi H, Bai M, Liu Y, Gao D, Tu D, Bao Y, Wang L, Chen S, Zhao X, Qu G. Genome sequence of Kobresia littledalei, the first chromosome-level genome in the family Cyperaceae. Sci Data. 2020; 7(1):175. doi: 10.1038/s41597-020-0518-3.

[48] Nong W, Qu Z, Li Y, Barton-Owen T, Wong AYP, Yip HY, Lee HT, Narayana S, Baril T, Swale T, et al. Horseshoe crab genomes reveal the evolution of genes and microRNAs after three rounds of whole genome duplication. Commun Biol. 2021; 4(1):83. doi: 10.1038/s42003-020-01637-2.

[49] Peterson KJ, Beavan A, Chabot PJ, McPeek MA, Pisani D, Fromm B, Simakov O. MicroRNAs as Indicators into the Causes and Consequences of Whole-Genome Duplication Events. Mol Biol Evol. 2022; 39(1):msab344. doi: 10.1093/molbev/msab344.

